# Integration of multi-source gene interaction networks and omics data with graph attention networks to identify novel disease genes

**DOI:** 10.1101/2023.12.03.569371

**Authors:** Kaiyuan Yang, Jiabei Cheng, Shenghao Cao, Xiaoyong Pan, Hong-Bin Shen, Cheng Jin, Ye Yuan

## Abstract

The pathogenesis of diseases is closely associated with genes, and the discovery of disease genes holds significant importance for understanding disease mechanisms and designing targeted therapeutics. However, biological validation of all genes for diseases is expensive and challenging. In this study, we propose DGP-AMIO, a computational method based on graph attention networks, to rank all unknown genes and identify potential novel disease genes by integrating multi-omics and gene interaction networks from multiple data sources. DGP-AMIO outperforms other methods significantly on 20 disease datasets, with an average AUROC and AUPR exceeding 0.9. The superior performance of DGP-AMIO is attributed to the integration of multiomics and gene interaction networks from multiple databases, as well as triGAT, a proposed GAT-based method that enables precise identification of disease genes in directed gene networks. Enrichment analysis conducted on the top 100 genes predicted by DGP-AMIO and literature research revealed that a majority of enriched GO terms, KEGG pathways and top genes were associated with diseases supported by relevant studies. We believe that our method can serve as an effective tool for identifying disease genes and guiding subsequent experimental validation efforts.

## Introduction

Researches have indicated a close association between diseases and genes^1-4^, and the existing catalog of disease genes is considered incomplete. Therefore, the discovery of disease genes holds profound significance as it contributes to our understanding of disease mechanisms and facilitates the design of targeted therapeutics^5^. However, the wet-lab experiments required to validate gene-disease associations are time-consuming and expensive. Moreover, considering the vast number of human genes, exceeding twenty thousand, it is unfeasible to perform biological experiments for all genes. Hence, accurate and efficient computational tools need to be developed to prioritize candidate genes and select those most likely to be associated with diseases. This would guide subsequent experimental validation and enhance the overall efficiency of the entire process.

The role of genes in diseases is diverse and complex. Some genes undergo mutations directly associated with diseases^3^, while others contribute to disease occurrence through abnormal expression^4^. Additionally, research has shown an association between DNA methylation and diseases^6-8^. Moreover, genes interact in intricate ways, including signal transduction, regulatory pathways, and protein-protein interactions. Some genes exert their effects on diseases by regulating or interacting with other genes^2, 9, 10^. For example, RAD51 is involved in recombination and DNA repair, and its expression products will form a complex with BRCA1 and BRCA2 proteins to participate in the process of DNA double-strand damage repair, which is related to the pathogenesis of breast cancer^2^. Inspired by these insights, numerous methods have been proposed for identifying disease genes. One category is based on epigenetics, where machine learning models are trained using omics data such as gene expression data to identify disease genes^11, 12^. Another category involves constructing gene interaction networks, such as protein-protein interaction networks, and using traditional graph machine learning methods to extract topological features of nodes to identify disease genes^13-16^. These aforementioned methods only utilize a single type of data, either multidimensional omics data or gene interactions. Considering the complexity of the gene-disease associations, integrating multiple types of data becomes necessary. The advancement of graph neural network (GNN) techniques provides a viable framework for this integration, where genes serve as nodes, gene interactions as edges, and multidimensional omics data as node features. This led to the emergence of GNN-based methods^17-21^. EMOGI^17^, for instance, combines protein-protein interaction networks with four types of omics data using graph convolutional network^22^ (GCN) to identify cancer genes, demonstrating the significant advantages of integrating gene interaction networks and omics data over using a single data type alone.

Nevertheless, there are some limitations in existing graph neural network-based methods. Firstly, the majority of these methods construct undirected graphs based on protein-protein interaction (PPI) networks. However, gene regulations and signaling pathways are directional, which are valuable sources of knowledge for predicting disease genes. Therefore, it is imperative to develop an effective approach that can leverage the directional information present in digraphs to enhance the accuracy of node classification tasks. Furthermore, there are diverse types of gene interaction networks available, including PPI networks, gene regulatory networks, and KEGG^23^ pathways, each originating from various databases. Existing methods often require training on separate networks from different databases. For instance, EMOGI^17^ are trained on six PPI networks individually. However, the performance of these models exhibits considerable variations across different databases, leading to disparate predictions for the same gene. Consequently, it is necessary to integrate gene interaction networks from multiple databases, constructing a more comprehensive knowledge graph and enabling more precise predictions of disease genes.

To address the issues mentioned above, we proposed a **D**isease **G**ene **P**redictor based on **A**ttention **M**echanism and **I**ntegration of multi-source gene interaction networks and **O**mics (DGP-AMIO). We merged gene interaction networks of different types and databases into a unified directed graph. Moreover, we introduced a 0/1 vector on the edges to indicate the presence or absence of gene interactions in each database and incorporated this edge feature into the training of attention coefficients. Additionally, we proposed triGAT improved based on graph attention networks^24^ (GAT), which better utilizes the relationships between genes and their upstream/downstream genes for disease gene prediction. Experimental results demonstrate that DGP-AMIO outperforms other methods on multiple diseases, achieving an average AUROC and AUPR above 0.9 on the test set. Ablation experiments show that both improvement strategies have a positive impact on the model performance. In addition, we conducted model behavior analysis, which elucidated DGP-AMIO’s ability to effectively integrate the topological features of the graph with omics features for accurate prediction of disease genes. Furthermore, gene enrichment analysis and literature review provide ample support evidence for the predictions made by DGP-AMIO. Overall, DGP-AMIO is a general computational tool capable of accurately predicting novel disease genes. The predictions by DGP-AMIO have valuable implications for guiding subsequent experimental validations.

## Results

DGP-AMIO is based on GATs and multi-graph integration, and trained in a semi-supervised framework to identify potential disease genes. It constructs a graph from gene interactions of multiple databases and gene expression profiles (Fig. 1a). In detail, gene regulatory networks and PPIs were downloaded from publicly available databases and converted into digraphs respectively, where nodes are genes and edges are interactions between them, and then all nodes and directed edges were taken union. A 0/1 vector was added on each edge to represent the database information, where 1 indicates that the gene interaction was recorded in the database of the vector’s corresponding dimension and 0 indicates unrecorded. Preprocessed gene expression or other omics data was combined with the graph as node feature. The graph was partially labelled, where positive labels are known disease genes recorded in Malacards and negative labels are genes randomly sampled from the remaining genes (Fig. 1b). To fully extract the upstream and downstream features of genes, we developed triGAT (Fig. 1c), a GAT-based model which can leverage direction information of edges in the digraph and learn better node representations. The graph was fed into triGAT, and the model was trained on five-fold cross validation dataset and tested on labeled data that did not appear in the training and validating process. The output of DGP-AMIO is the probabilities of being disease genes on all unlabeled data in the graph. (Fig. 1d)

**Fig. 1.**
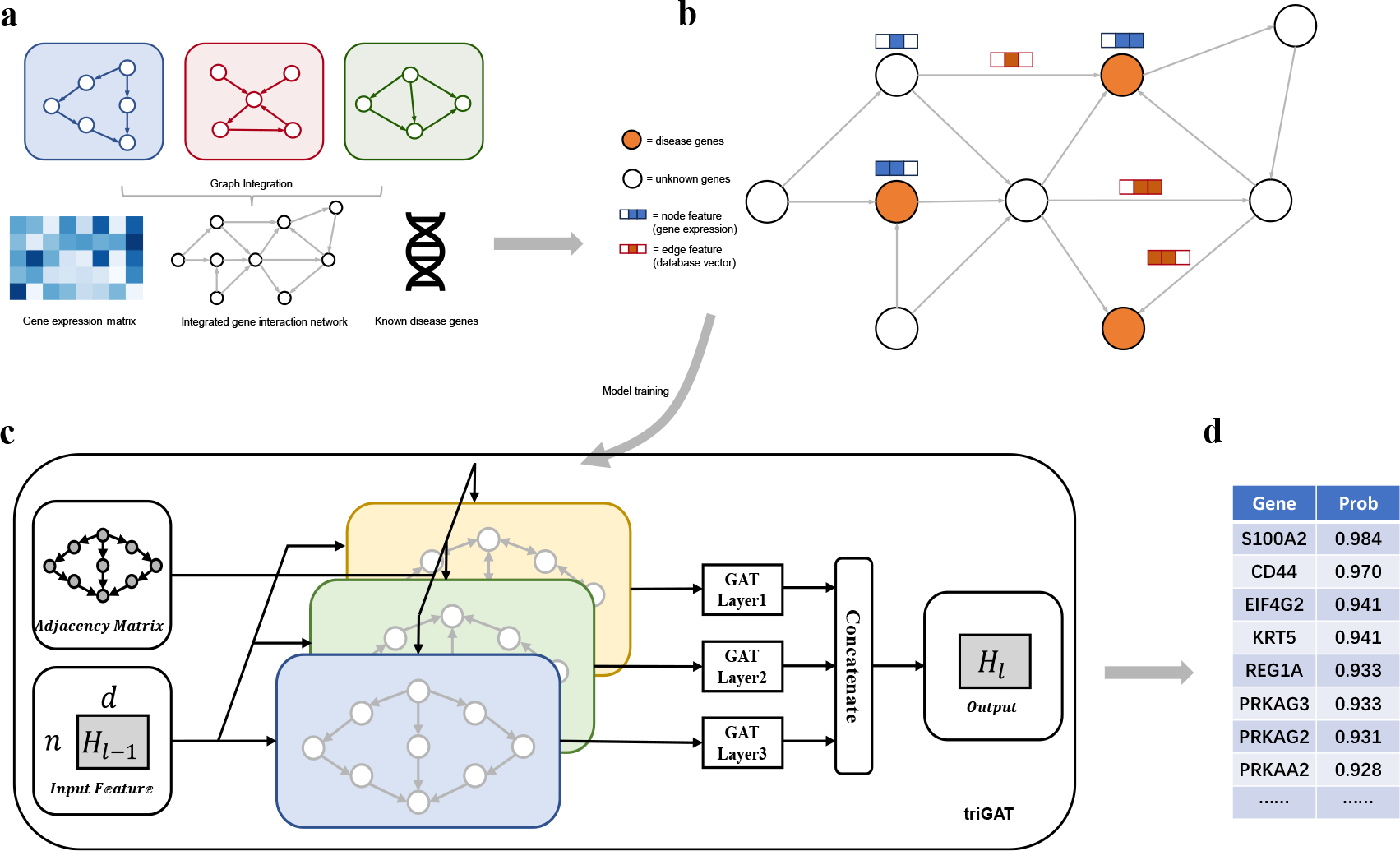
Schematic of DGP-AMIO framework. **a**, Data preparation and integration. Gene interaction networks from different databases are integrated to form a unified digraph which is then combined with gene expression (or other omics) data and knowledge of known disease genes. **b**, The knowledge graph for training, where nodes correspond to genes, edges to interactions, node features to multidimensional gene expression vectors and edge features to the records of gene interactions in different databases, which are d-dimension 0/1 vectors. (d is the number of databases). **c**, framework of triGAT, proposed in DGP-AMIO to better extract node embeddings in directed graph for node classification. triGAT takes as input the digraphs and is trained on the original graph, reverse graph and bidirectional graph and concatenated the node embeddings of the three graphs to obtain the output. **d**, The output of DGP-AMIO, a gene list ranked according to the probability of being related to diseases.

### DGP-AMIO precisely identifies disease genes

We trained DGP-AMIO on disease genes from Malacards based on expression data from Gene Expression Omnibus (GEO) and the integration of ten gene interaction networks from publicly available databases. We evaluated DGP-AMIO’s performance across different diseases.

#### Comparison with other methods

We compared DGP-AMIO with other methods for disease gene prediction on 20 diseases, including cancer and non-cancer diseases. For each disease, we divided the dataset into the training and test set and conducted 5-fold cross validation on the training set, and computed the AUROC and AUPR on the test set for each method.

To demonstrate the superiority of our method, we benchmarked DGP-AMIO against three methods. CNNC^11^ is a deep learning method for inferring gene relationships from expression data based on CNN, which converted the coexpression data of gene pairs into images and fed them into CNN. It can predict new disease genes through their coexpression with known disease genes. EMOGI^17^ is a pan-cancer gene identification method based on GCN, using multiomics data and PPI networks as input, which can also be used for disease gene prediction. Furthermore, we benchmarked DGP-AMIO against node2vec+RandomForest, which also combines omics data and graph structure information. This method was shown to have similar performance to EMOGI. The graph features learned by node2vec was concatenated with gene expression data and then were fed into a random forest binary classifier. For fair comparison, we used the same gene expression data for all methods and used STRING PPI network for EMOGI and node2vec+RandomForest. Moreover, to prove DGP-AMIO performs better in predicting novel disease genes, we compared the four methods on disease genes not included in training and test set. We collected disease related genes mainly from DisGeNet^25^, a database of gene-disease associations from expert curated repositories, GWAS catalogues, animal models and the literature, and for asthma we also collected disease genes from AllerGAtlas^26^, a manually curated human allergy-related gene database. Then, we removed genes that appear in the training and test set. The remaining genes were taken as positive samples and an equal number of genes were randomly sampled from the other genes as negative samples to calculate AUROC and AUPR.

The results show that the performance of DGP-AMIO significantly outperforms other methods across all diseases, with AUROC and AUPR above 0.9 on the test set for most diseases (Fig. 2a). Fig. 2c shows the results of five-fold cross validation on six diseases. Additionally, DGP-AMIO also outperforms other methods in predicting novel disease genes not appearing in the training and test set (Fig. 2b), demonstrating the superior potential of DGP-AMIO in discovering new disease genes.

**Fig. 2.**
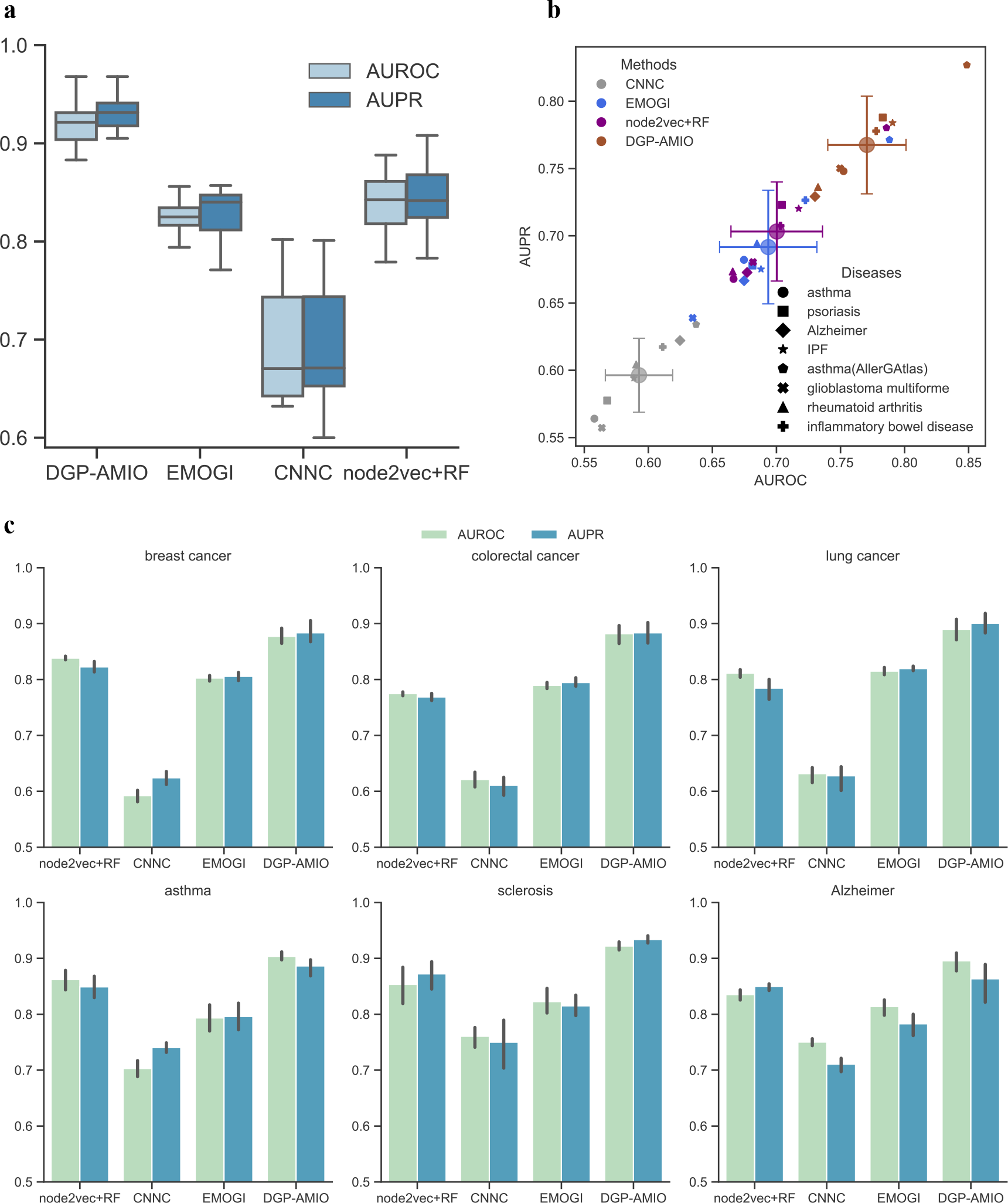
DGP-AMIO outperforms previous methods in predicting disease genes on multiple diseases. **a**, AUROCs and AUPRs of different methods, computed on the test sets of 20 diseases. **b**, Performance comparison of different methods on genes not included in the training and test set, collected from other datasets, DisGeNet and AllerGAtlas. The large circles correspond to average AUROCs and AUPRs across diseases for each method, and the crosses represent variances. **c**, Five-fold cross validations of different methods on six diseases: breast cancer, colorectal cancer, lung cancer, asthma, sclerosis and Alzheimer.

#### triGAT improves DGP-AMIO’s capability to identify both upstream and downstream disease genes

The digraph integrating directional gene regulatory relations contains more information than the undirected graph, which can be viewed as the weakened version of the digraph. We expect the model to integrate the direction information for accurate prediction. To prove that triGAT has better ability to extract structural features of digraph and predict disease genes, we compared triGAT with GAT and GCN based on the same graph. The input graph of triGAT and GAT was exactly the same while the input of GCN was the undirected graph derived from the original one. The AUROC and AUPR on the test set shows that triGAT is the best GNN model for predicting disease genes, followed by GCN and GAT (Fig. 3a). Although GAT receives the digraph as input, it yields the worst performance. We examined the disease-related probability of the disease genes in the test set predicted by GAT and found that the genes located downstream of known disease genes had higher probabilities than upstream disease genes (Fig. 3c, d). This phenomenon can be explained by the message passing mechanism in GAT, where message can only flow from upstream node to downstream node. GCN does not have this problem, which results in better performance than GAT, but also leads to a lack of directional information. triGAT enables messages to be passed from downstream to upstream while retaining the original digraph structure. Fig. 3b-d shows that the upstream gene prediction accuracy of triGAT is significantly better than GAT and the downstream genes can also be more accurately predicted.

**Fig. 3.**
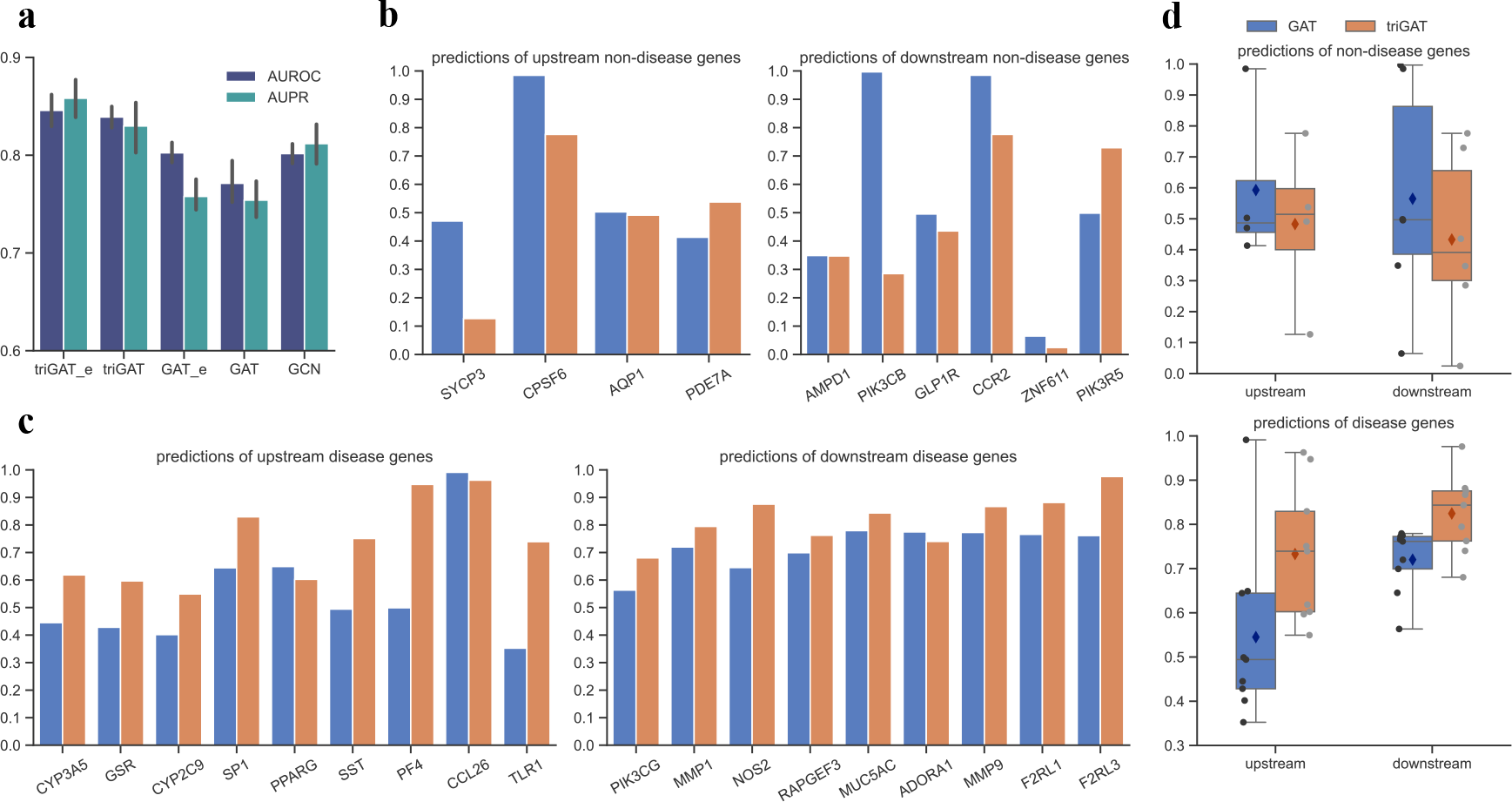
triGAT enables DGP-AMIO to precisely identify upstream and downstream disease genes. **a**, Performance comparison of triGAT, GAT and GCN. ‘_e’ means edge features are included. **b-d**, Predictions of triGAT and GAT on non-disease genes (**b**) and disease genes (**c**) in the test set which are located upstream and downstream of the disease genes in the training set, and the statistical distribution (**d**), where the diamond points represent mean values.

#### DGP-AMIO benefits from the integration of multiple gene interaction databases and multiomics

To evaluate whether DGP-AMIO trained on the graph integrating multiple databases was better than DGP-AMIO trained on the graph of a single database. We conducted experiments for breast cancer, given its high number of known disease genes available for training. We first trained DGP-AMIO on six single gene interaction networks. The AUROC and AUPR of DGP-AMIO fluctuated significantly with different single database (Fig. 4a), which suggests that for graph-based method, the accuracy of predicting disease genes is considerably affected by the gene interaction network used. Next, we trained DGP-AMIO on the integrated network. The results showed that as the number of integrated gene interaction databases increased (from 1 to 10), the AUROC and AUPR of DGP-AMIO increased and is significantly higher than DGP-AMIO trained on any single database (Fig. 4b), which proved the effectiveness of multi-database integration. Furthermore, we trained DGP-AMIOs with and without database vectors on edges and compared their performances. Fig. 3a shows that adding vectors containing database information on edges improved AUROC and AUPR of DGP-AMIO. This demonstrates that incorporating edge features into the training of attention coefficients enables the DGP-AMIO to learn the importance of different gene interaction data and allocate attention weights based on the database record.

**Fig. 4.**
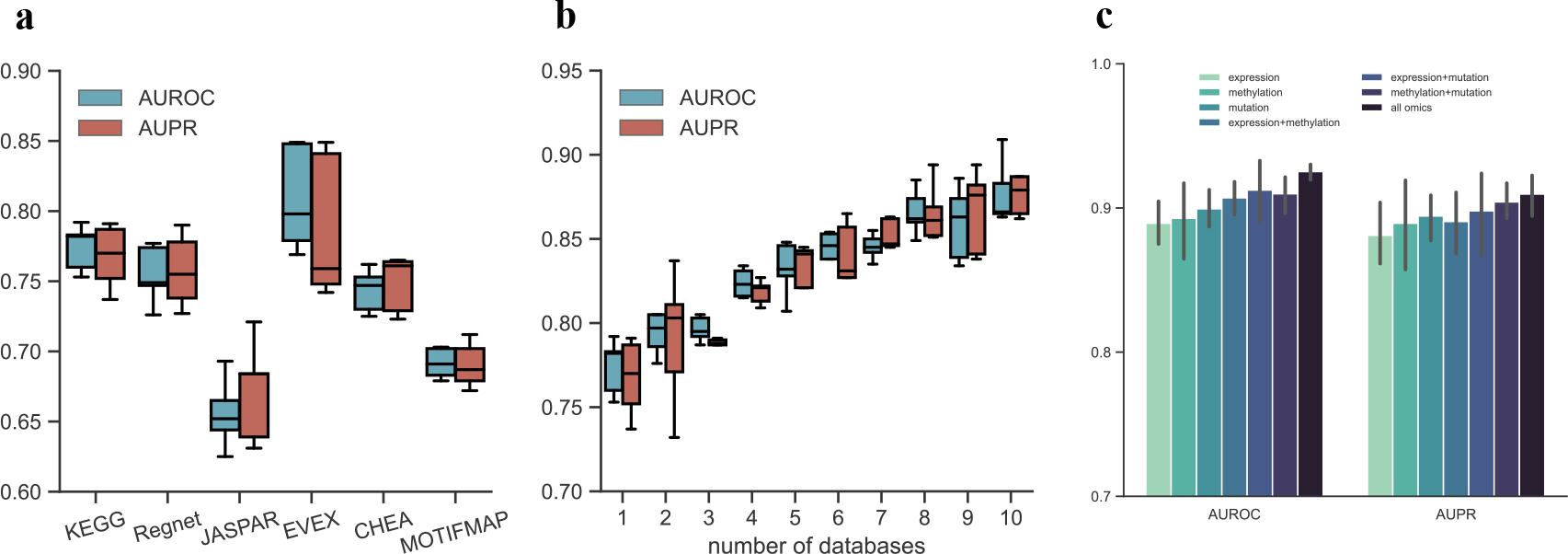
DGP-AMIO benefits from the integration of gene interaction databases and multiomics. **a**, Performances of DGP-AMIO trained on different single gene interaction network. **b**, Performances of DGP-AMIO integrating different number of gene interaction networks, from 1 to 10. **c**, Performances of DGP-AMIO trained using one type of omics, two types and all three types (expression, DNA methylation and gene mutation).

Moreover, DGP-AMIO is a general framework that can integrate multiomics data by simply concatenating multiple omics as node features. To explore whether multiomics integration can improve the performance of DGP-AMIO, we downloaded from TCGA the expression, DNA methylation and gene mutation data, which are associated with diseases^6-8, 27^, and trained DGP-AMIO using single type of omics, two types of omics and all three omics and evaluate their performances. Fig. 4c is the result of colorectal cancer, showing that the AUROC and AUPR of DGP-AMIO using three omics is the highest, and using two omics is better than single omics. This demonstrates the multiomics integration can further improve DGP-AMIO’s ability to predict disease genes.

### Model behavior analysis

To comprehend the predictions of DGP-AMIO, we conducted model behavior analysis focusing on the topological characteristics of graphs and differential expression analysis, using prostate cancer as example.

We calculated the pearson correlation coefficients (PCC) between the DGP-AMIO score (the probability of a gene of being a disease gene) and the number of disease neighbors (PCC=0.63), disease predecessors (PCC=0.57) and disease successors (PCC=0.61) of that gene (Fig. 5a) and found a significant correlation between them. This indicates the potential disease genes predicted by DGP-AMIO exhibit close interactions with existing disease genes. Additionally, experimental evidence demonstrated that incorporating omics data, such as gene expression, as node features leads to improved predictive performance compared to training solely based on the graph structure (Fig. 5c), hence we further performed differential expression analysis of GSE114740 (Fig. 5b), mRNA profiles of 10 pairs of localized primary prostate cancer samples and the matched normal tissues.

**Fig. 5.**
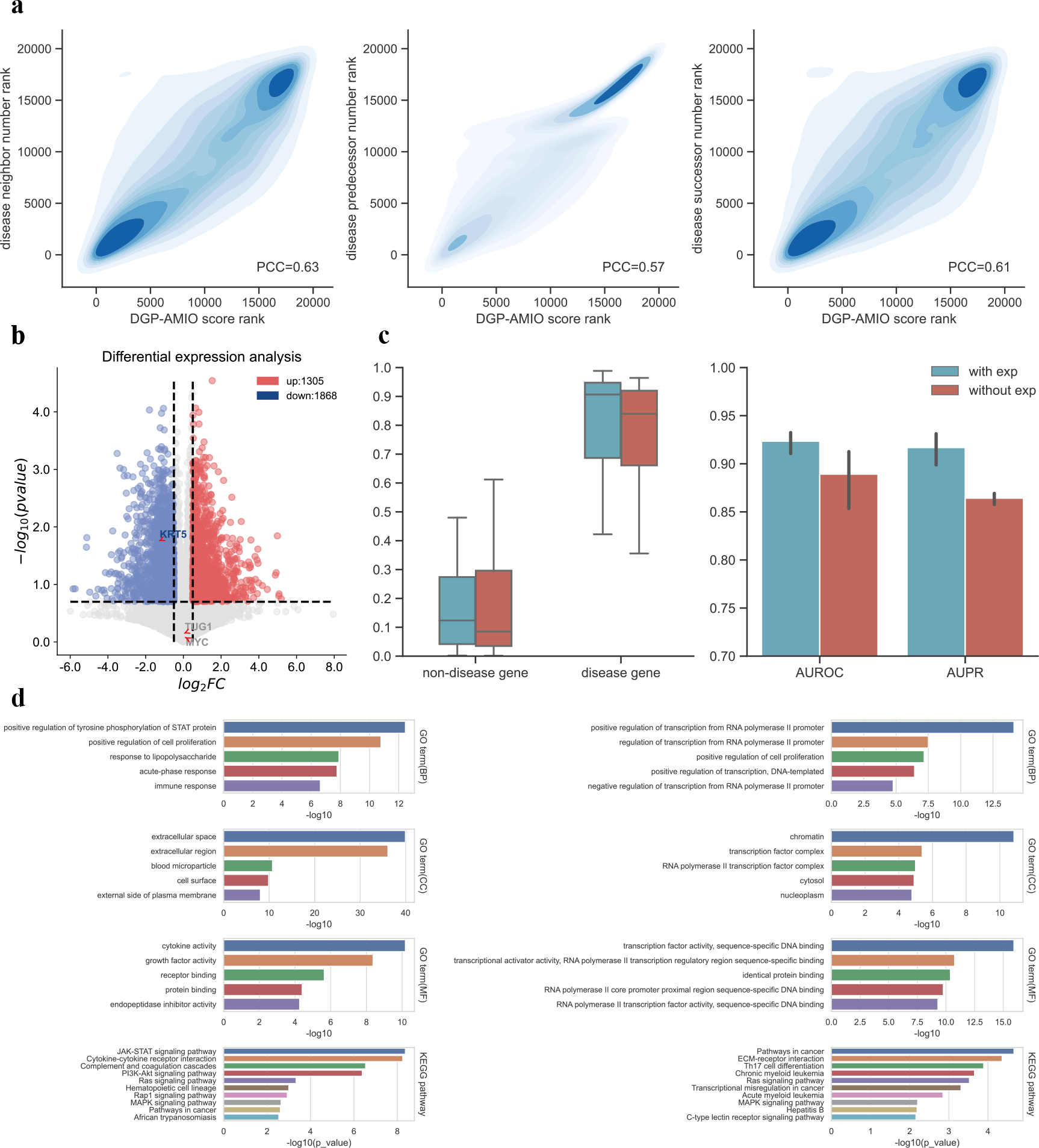
Model behavior analysis and enrichment analysis. **a**, rank correlation plots between DGP-AMIO score (x-axis) and the number of disease neighbors, predecessors and successors (y-axis). Genes with high probability of being disease genes tend to have numerous interactions with known disease genes. **b**, differential expression analysis of GSE114740. **c**, performance comparison between DGP-AMIO trained on graph only and graph with gene expression as node feature. **d**, enrichment analysis of top 100 genes predicted by DGP-AMIO (left: asthma, right: colorectal cancer), showing top 5 enriched GO terms of biological process (BP), cellular component (CC), molecular function (MF) and top 10 enriched KEGG pathways and their p values.

We selected MYC, KRT5 and TUG1 gene for analysis. They are all known prostate cancer genes in the test set and accurately predicted with top 20 DGP-AMIO score among all genes except genes in the training set. MYC does not exhibit significant expression differences between cancer samples and normal tissues. However, it interacts with 420 prostate cancer genes in the graph, ranking third among all genes. KRT5 interacts with 36 prostate cancer genes in the graph, ranking 1961 among all genes, while it exhibits significant expression differences. For TUG1, it exhibits neither significant expression differences nor extensive interactions with prostate cancer genes (only interacts with ETS2 and YY1). However, the two genes it interacts with, ETS2 and YY1, are crucial hub cancer genes interacting with 377 and 473 prostate cancer genes, which are second-order neighbors of TUG1. The aforementioned analysis indicates that graph structure and omics complement each other, as their combination provides more comprehensively informative signals for learning by DGP-AMIO, leading to more accurate predictions. Furthermore, the multi-layer graph neural network architecture can capture high-order neighborhood information in the graph, further enhancing the prediction accuracy.

### Newly predicted disease genes

#### Enrichment analysis

To further evaluate the performance of our method, we performed gene enrichment analysis on the top 100 genes of DGP-AMIO’s predictions. Fig. 5 shows the result of asthma and colorectal cancer, listing top 5 GO terms of biological process (BP), cellular component (CC), molecular function (MF) and top 10 enriched KEGG pathways and the corresponding p values.

For asthma, the most significantly enriched GO terms are “positive regulation of tyrosine phosphorylation of STAT protein”, “positive regulation of cell proliferation”, “extracellular space”, “extracellular region”, “cytokine activity” and “growth factor activity”, all of which are related to asthma. Protein tyrosine phosphorylation of STAT is involved in regulating the immune response and its inhibition can reduce the development of asthma^28^. The cell proliferation of airway smooth muscle cells stimulated by airway inflammation is recognized to be crucial to asthma^29, 30^. “Extracellular space” and “extracellular region” are asthma-related because extracellular vesicles, matrix components and DNA play an important role in the pathogenesis of asthma^31-33^. Cytokine activity is closely associated with asthma, as cytokines like IL-4, IL-5, and IL-13 can promote mucus overproduction and bronchial hyperresponsiveness^34^, which are symptoms of asthma. Many growth factors are also involved in asthma such as EGF, FGF and TGFs^35^. Among the enriched KEGG pathways, “JAK-STAT signaling pathway”, “cytokine-cytokine receptor interaction”, “complement and coagulation cascades” and “PI3K-Akt signaling pathway” have the lowest p value. JAK-STAT signaling pathway transduces cytokine-mediated signals which are pivotal in the development of asthma^36^, and targeting this pathway has been verified to have a therapeutic effect on asthma^37^. Cytokine-cytokine receptor interaction is enriched because of the vital role of cytokines in asthma discussed above. Complement and coagulation system is found to be in association with asthma control^38^. The PI3K-Akt signaling pathway can promote airway inflammation and hyperresponsiveness^39^, therefore is crucial in asthma.

For colorectal cancer, the top enriched GO terms are “regulation of transcription from RNA polymerase II”, “positive regulation of cell proliferation”, “chromatin”, “transcription factor complex”, “transcription factor activity” and “transcription activator activity”, which are all cancer-related. RNA polymerase II is related to disruption of transcription elongation which is implicated in cancer^40^. The abnormal cell proliferation is the main cause of various cancer. “Chromatin” is cancer-related as its dysregulation is linked to mutation and cancer^41^. Cancer development requires constitutive expression/activation of transcription factors (TFs) and activators for growth and survival^42^ and many TFs are tumor suppressors or oncogenes. The most significantly enriched KEGG pathways are “pathways in cancer” (evidently cancer-related), “ECM-receptor interaction” and “Th17 cell differentiation”. The ECM receptors constitute crucial pathways involved in colorectal cancer progression and metastasis^43^ and Th17 cells are also highly related to colorectal cancer progression.

In conclusion, the top genes predicted by DGP-AMIO are significantly enriched in disease-related GO terms and pathways, demonstrating DGP-AMIO’s ability of discovering novel disease genes.

#### Relevant research about the predicted unknown genes

We listed the top 10 potential disease genes of asthma and predicted by DGP-AMIO, as shown in Table. 1. We searched them online and found that most of them were studied as disease-related genes in the previous research, which indicates the prediction of DGP-AMIO is consistent with the existing studies. This demonstrates that DGP-AMIO is an effective computational method for discovering novel disease genes.

**Table 1.**
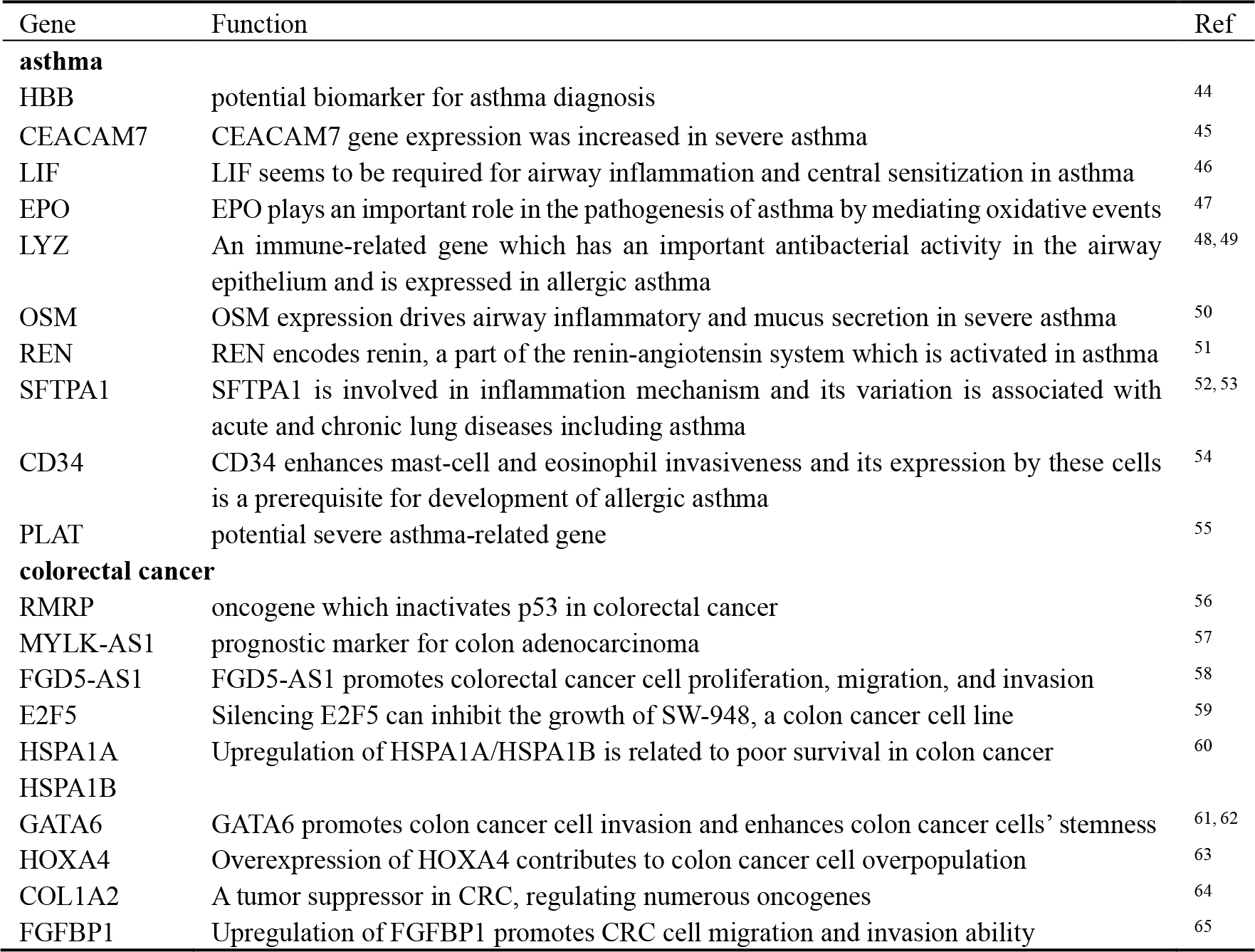
Top 10 ranked genes of asthma and colorectal cancer predicted by DGP-AMIO and their functions in disease supported by relevant studies.

## Discussion

In this work we proposed DGP-AMIO, a GAT-based disease gene predictor with integration of gene interaction networks from different databases and multiomics. We demonstrated DGP-AMIO has a higher accuracy and generalization than existing methods across 20 disease datasets, with AUROC and AUPR of most diseases reaching above 0.9 on test set, and DGP-AMIO also outperforms other methods on unlabeled genes during the training and testing period. Additionally, we performed model behavior analysis and concluded that DGP-AMIO can effectively integrate the topological feature of graph, high-order neighborhood information and omics feature for accurate prediction. Furthermore, we conducted enrichment analysis and literature search on the top genes predicted by DGP-AMIO, with results showing most enriched GO terms/pathways and top genes are disease-related supported by existing literature. This provides compelling evidence for the efficacy and accuracy of our method as a tool for predicting novel disease-associated genes.

We concluded several improvements of DGP-AMIO compared to existing methods. First, DGP-AMIO was trained on a digraph and triGAT was proposed to extract better node embedding from a digraph. Experiments showed the retainment of direction of gene interactions contributed to DGP-AMIO’s better performance. Second, DGP-AMIO was trained on an integrated graph including multiple databases of gene regulatory networks and PPI networks. This not only signifies the utilization of a broader range of knowledge information, but also eliminates the need for training separate graphs on different databases, which may result in variations in the predicted outcomes across different databases. This integration significantly enhances the performance of DGP-AMIO in predicting disease genes. Finally, DGP-AMIO is a general framework which can integrate multiomics and multiple gene interaction networks, and it can be applied for disease gene prediction for a wide range of diseases, as long as disease-related omics data is provided.

We acknowledge several limitations of DGP-AMIO. We employed random sampling from non-known disease genes as negative samples for training. Experimental observations revealed that different sampling can influence DGP-AMIO’s performance, highlighting the need for improved negative sample construction strategies. The quality of omics data used as input can also affect the performance. Although the integration of multiomics has demonstrated improvement for certain diseases, this may not be applicable to all diseases. Additionally, DGP-AMIO exhibits limited predictive performance for diseases with a scarce number of known disease genes documented in Malacards. Regarding the aforementioned issues, future improvement work will focus on the following aspects: 1. Developing more appropriate strategies for constructing negative samples, including but not limited to, introducing constraints during sampling and incorporating information from relevant studies. 2. Utilizing gene embeddings obtained from large models pretrained on large-scale transcriptomic data. 3. Augmenting the scale of labeled samples by integrating literature data and clinical trial data. In summary, DGP-AMIO is a general computational approach capable of accurately predicting disease-associated genes, providing a powerful tool for the discovery of novel disease genes.

## Methods

### Data acquisition and preprocessing

#### Gene expression data

We mainly collected gene expression data from GEO, and for cancer gene prediction, TCGA is also an available database providing different types of omics data. We averaged duplicate gene expression and deleted genes whose expression quantities of all samples are zero. Expression data only is enough for model training, we can use other types of omics such as DNA methylation or gene mutation and concatenate them all for better performance.

#### Integration of multiple gene interaction networks

We collected gene regulatory networks including KEGG^23^, RegNetwork^66^, TRRUST^67^, HTRIdb^68^, EVEX^69^, JASPAR^70^, CHEA^71^, TRANSFAC^72^, MOTIFMAP^73^, and ENCODE^74^, and PPI networks including CPDB^75^, STRING^76^, IRefIndex^77^, and PCNet^78^, for network integration. Depending on the networks, we preprocessed the data with different methods. For KEGG, we downloaded 345 pathways of human through KEGGREST, converted them into directed graphs and merged them through KEGGgraph in R, generating a directed graph including all pathways. For CPDB, STRING and RegNetwork, which contain confidence information of gene-gene interactions, we retained interactions with high confidence. Referring to the processing method in the work of EMOGI^17^, we retained interactions with a score higher than 0.5 for CPDB and higher than 0.85 for STRING, and for RegNetwork we retained gene regulations whose confidence is high or middle. For the IRefIndex, we only retained binary interactions. Networks from other databases were directly downloaded without further preprocessing.

Before the integration of multiple databases, considering the PPI networks are undirected graph while our model deals with directed graph, we converted each undirected edge in PPI networks into two reverse directed edges. Then we take the union of edges in all databases. For each edge, we used a d-dimensional vector composed of 0 and 1 to represent its database information, where d equals the number of databases and 1 means that this interaction is recorded in the database of the corresponding dimension. In the model this vector appears as edge feature and is involved in training the attention weights.

#### Disease genes

We collected known disease genes from Malacards, an integrated database of human maladies. These genes are positive samples with label 1 in model training.

#### Data Overlap

For genes without expression (or other omics) data, we deleted the corresponding nodes in the network and their connected edges. Moreover, we removed the known disease genes not covered in the network. In this way we constructed a knowledge graph, where nodes and edges are genes and interactions, gene expression data are node feature, database information is edge feature, and genes with label 1 are disease genes.

### Model

#### GATs

Our model structure is based on the graph attention networks (GAT)^24^. GAT is a spatial graph convolutional network based on attention mechanism. During message passing, instead of simply summing or averaging neighbor node features, GAT assigns different importance to neighbor nodes through computing attention coefficient:

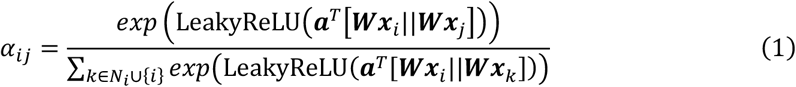

The attention coefficient *α*_*i*_ represents the importance of node *j* to node *i*, where *x*_*i*_, *x*_*i*_ ∈ *R*^*F*^ are node feature, 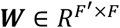 is a trainable weight matrix, 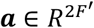 is a weight vector representing a shared single-layer feedforward neural network, and *N*_*i*_ is the set of neighbor nodes of node *i*. In directed graph where information flows in the direction of edges, *N*_*i*_ is the set of neighbor nodes pointing to *i*. Then the model aggregates message from neighbor nodes by computing weighted sum and applying a nonlinearity σ, hence generates the updated representation of node *i*:

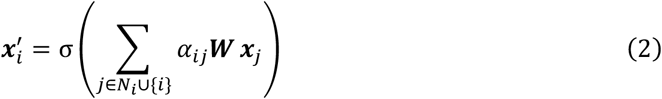

Moreover, GAT applies multi-head attention mechanism, allowing that different attention coefficients are computed between the same nodes by learning ***a***^*k*^(the *k*-th attention mechanism represented by a single-layer FFN, *k* = 1, …, *K*), so that GAT can capture richer information. For *K*-head attention, the output node representation of one GAT layer is computed by concatenating or averaging the *K* updated node features:

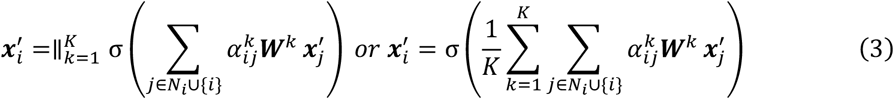

where 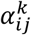 is the attention coefficient computed by ***a***^*k*^, and ***W***^*k*^ is the corresponding linear transformation weight matrix, both of which are trainable.

#### triGATs

##### Attention mechanism with edge feature

Considering our model includes edge features which represent the database information, we added them into the calculation of attention coefficients:

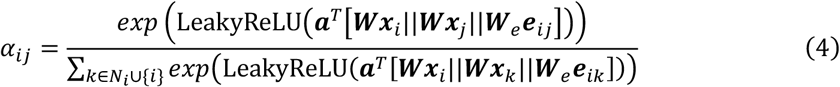

where ***e***_*ij*_ is the feature of edge *j* → *i*, ***W***_***e***_ is the transformation weight matrix of edge feature. This allows GAT to automatically learn the importance of different databases.

##### triGAT with message feedback

When GAT handles directed graphs, the message can only flows from the source node to the target node, while cannot be fedback from downstream to upstream. One feasible solution is to convert all directed edges to undirected ones, while this may cause the loss of directional information, which is critical in gene regulatory networks and pathways. To address this problem, we proposed triGAT, which not only retains the directional information in the original digraph, but also enables message feedback from downstream to upstream. One triGAT layer is composed of three single-layer GATs whose input graphs are *G*_1_, *G*_2_ and *G*_3_. *G*_1_ is the original digraph, *G*_2_ is the converse digraph where direction of each edge in *G*_2_ is opposite to *G*_1_, and *G*_3_ is the bidirectional graph where between each connected node pair exists two opposite edges. The input node and edge features of these three GATs are the same. triGAT layer concatenates the learned node representations of three GATs and propagates them forward to the next layer, as shown in equation (5), where ***X, E*** are the input node and edge feature and ***X***^′^ is the output node representation matrix of one triGAT layer.

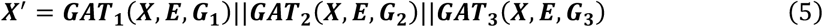

### Model Training

In disease gene prediction tasks, known disease targets (recorded in Malacards) are positive samples while the other gene labels are unknown, which means we lack samples with clear negative label. Considering minimal genes are related to human diseases, we randomly sampled genes equal to the number of positive samples among the unlabeled genes as negative samples.

After sampling, the labelled data were first randomly split into the training (80%) and test (20%) set, and then the training set was divided into five equal parts for cross validation, with each split keeping the ratio of positive and negative samples unchanged. During training, the training set was used to train five ensembles, each ensemble holding out one part for validation and early stopping. The input of the model was composed of a node feature matrix ***X***, an integrated gene interaction network represented by its adjacency matrix *A* and edge feature matrix *E*, and labels *y*. We applied cross-entropy as loss function and used ADAM as optimizer to train our model. The earlystopping strategy was used to stop training when the loss on validation set failed to decrease for *p* consecutive epochs.

### Evaluation and Prediction

The average output of five ensemble models generated by cross validation was used for calculation of AUROC and AUPR on test set. To further evaluate the precision and generalization, we collected disease-associated genes from DisGeNet and AllerGAtlas and removed the genes that appeared in the training and test set. Then we used the remaining genes as positive and sampled unknown genes as negative to calculate metrics. After evaluation, the model can predict on the whole graph, giving a disease-related probability value for each gene except for known disease genes. We sorted these genes in descending order of the probability and the top genes are more likely to be novel disease targets.

## Code availability

The code to train DGP-AMIO, evaluate model performance and predict disease genes is available at https://github.com/yangkaiyuan1027/DGP-AMIO.

## References

1. Jackson, M., Marks, L., May, G.H. & Wilson, J.B. The genetic basis of disease. Essays in biochemistry 62, 643–723 (2018).

2. Kato, M. et al. Identification of Rad51 alteration in patients with bilateral breast cancer. J Hum Genet 45, 133–137 (2000).

3. Machackova, E. et al. Spectrum and characterisation of BRCA1 and BRCA2 deleterious mutations in high-risk Czech patients with breast and/or ovarian. Bmc Cancer 8 (2008).

4. Zheng, S.P. et al. Overexpression of CBX2 in breast cancer promotes tumor progression through the PI3K/AKT signaling pathway. Am J Transl Res 11, 1668–1682 (2019).

5. Linehan, W.M. et al. Identification of the genes for kidney cancer: Opportunity for disease-specific targeted therapeutics. Clin Cancer Res 13, 671s–679s (2007).

6. Calle-Fabregat, C.D., Morante-Palacios, O. & Ballestar, E. Understanding the Relevance of DNA Methylation Changes in Immune Differentiation and Disease. Genes-Basel 11 (2020).

7. Greenberg, M.V.C. & Bourc’his, D. The diverse roles of DNA methylation in mammalian development and disease. Nat Rev Mol Cell Bio 20, 590–607 (2019).

8. Skvortsova, K., Stirzaker, C. & Taberlay, P. The DNA methylation landscape in cancer. Essays Biochem 63, 797–811 (2019).

9. Hu, X. et al. LSD1 suppresses invasion, migration and metastasis of luminal breast cancer cells via activation of GATA3 and repression of TRIM37 expression. Oncogene 38, 7017–7034 (2019).

10. Zhang, Y.M. et al. The ORMDL3 Asthma Gene Regulates and Has Multiple Effects on Cellular Inflammation. American Journal of Respiratory and Critical Care Medicine 199, 478–488 (2019).

11. Yuan, Y. & Bar-Joseph, Z. Deep learning for inferring gene relationships from single-cell expression data. Proceedings of the National Academy of Sciences 116, 27151–27158 (2019).

12. Xiao, Y. et al. Differential expression pattern-based prioritization of candidate genes through integrating disease-specific expression data. Genomics 98, 64–71 (2011).

13. Peng, J.J., Guan, J.J. & Shang, X.Q. Predicting Parkinson’s Disease Genes Based on Node2vec and Autoencoder. Front Genet 10 (2019).

14. Zhang, Y. et al. Identifying Breast Cancer-Related Genes Based on a Novel Computational Framework Involving KEGG Pathways and PPI Network Modularity. Front Genet 12 (2021).

15. Cowen, L., Ideker, T., Raphael, B.J. & Sharan, R. Network propagation: a universal amplifier of genetic associations. Nat Rev Genet 18, 551–562 (2017).

16. Liu, H.J. et al. Predicting the Disease Genes of Multiple Sclerosis Based on Network Representation Learning. Front Genet 11 (2020).

17. Schulte-Sasse, R., Budach, S., Hnisz, D. & Marsico, A. Integration of multiomics data with graph convolutional networks to identify new cancer genes and their associated molecular mechanisms. Nat Mach Intell 3, 513–526 (2021).

18. Han, P. et al. GCN-MF: Disease-Gene Association Identification By Graph Convolutional Networks and Matrix Factorization. Kdd’19: Proceedings of the 25th Acm Sigkdd International Conferencce on Knowledge Discovery and Data Mining, 705–713 (2019).

19. Zhang, T. et al. GCN-GENE: A novel method for prediction of coronary heart disease-related genes. Comput Biol Med 150 (2022).

20. Schulte-Sasse, R., Budach, S., Hnisz, D. & Marsico, A. Graph Convolutional Networks Improve the Prediction of Cancer Driver Genes. Lect Notes Comput Sc 11731, 658–668 (2019).

21. Azadifar, S. & Ahmadi, A. A novel candidate disease gene prioritization method using deep graph convolutional networks and semi-supervised learning. Bmc Bioinformatics 23 (2022).

22. Kipf, T.N. & Welling, M. Semi-supervised classification with graph convolutional networks. arXiv preprint arXiv:1609.02907 (2017).

23. Kanehisa, M. & Goto, S. KEGG: kyoto encyclopedia of genes and genomes. Nucleic acids research 28, 27–30 (2000).

24. Veličkovicć, P. et al. Graph attention networks. arXiv preprint arXiv:1710.10903 (2017).

25. Piñero, J. et al. DisGeNET: a comprehensive platform integrating information on human disease-associated genes and variants. Nucleic Acids Research 45, D833–D839 (2017).

26. Liu, J. et al. AllerGAtlas 1.0: a human allergy-related genes database. Database 2018, bay010 (2018).

27. Timar, J. & Kashofer, K. Molecular epidemiology and diagnostics of KRAS mutations in human cancer. Cancer Metast Rev 39, 1029–1038 (2020).

28. Pouliot, P., Bergeron, S., Marette, A. & Olivier, M. The role of protein tyrosine phosphatases in the regulation of allergic asthma: implication of TC-PTP and PTP-1B in the modulation of disease development. Immunology 128, 534–542 (2009).

29. Johnson, P.R. et al. Airway smooth muscle cell proliferation is increased in asthma. American journal of respiratory and critical care medicine 164, 474–477 (2001).

30. Khan, M.A. Inflammation signals airway smooth muscle cell proliferation in asthma pathogenesis. Multidisciplinary respiratory medicine 8, 1–5 (2013).

31. Alashkar Alhamwe, B. et al. Extracellular vesicles and asthma—More than just a co-existence. International journal of molecular sciences 22, 4984 (2021).

32. Araujo, B.B. et al. Extracellular matrix components and regulators in the airway smooth muscle in asthma. European Respiratory Journal 32, 61–69 (2008).

33. Lachowicz-Scroggins, M.E. et al. Extracellular DNA, neutrophil extracellular traps, and inflammasome activation in severe asthma. American journal of respiratory and critical care medicine 199, 1076–1085 (2019).

34. Lambrecht, B.N., Hammad, H. & Fahy, J.V. The cytokines of asthma. Immunity 50, 975–991 (2019).

35. Kardas, G. et al. Role of platelet-derived growth factor (PDGF) in asthma as an immunoregulatory factor mediating airway remodeling and possible pharmacological target. Frontiers in pharmacology 11, 47 (2020).

36. Pernis, A.B. & Rothman, P.B. JAK-STAT signaling in asthma. The Journal of clinical investigation 109, 1279–1283 (2002).

37. Vale, K. Targeting the JAK-STAT pathway in the treatment of ‘Th2-high’severe asthma. Future medicinal chemistry 8, 405–419 (2016).

38. Kokelj, S. et al. Activation of the Complement and Coagulation Systems in the Small Airways in Asthma. Respiration 102, 621–631 (2023).

39. Medina-Tato, D.A., Ward, S.G. & Watson, M.L. Phosphoinositide 3-kinase signalling in lung disease: leucocytes and beyond. Immunology 121, 448–461 (2007).

40. Cermakova, K. et al. A ubiquitous disordered protein interaction module orchestrates transcription elongation. Science 374, 1113-+ (2021).

41. Sehgal, P. & Chaturvedi, P. Chromatin and Cancer: Implications of Disrupted Chromatin Organization in Tumorigenesis and Its Diversification. Cancers 15, 466 (2023).

42. Vishnoi, K., Viswakarma, N., Rana, A. & Rana, B. Transcription Factors in Cancer Development and Therapy. Cancers 12 (2020).

43. Nersisyan, S. et al. ECM-Receptor Regulatory Network and Its Prognostic Role in Colorectal Cancer. Front Genet 12 (2021).

44. Zhou, Y. et al. Quantitative proteomics profiling of plasma from children with asthma. International Immunopharmacology 119, 110249 (2023).

45. Shikotra, A. et al. A CEACAM6-high airway neutrophil phenotype and CEACAM6-high epithelial cells are features of severe asthma. The Journal of Immunology 198, 3307–3317 (2017).

46. Lin, M.-J., Lao, X.-J., Liu, S.-M., Xu, Z.-H. & Zou, W.-F. Leukemia inhibitory factor in the neuroimmune communication pathways in allergic asthma. Neuroscience letters 563, 22–27 (2014).

47. Wu, Z., Mehrabi Nasab, E., Arora, P. & Athari, S.S. Study effect of probiotics and prebiotics on treatment of OVA-LPS-induced of allergic asthma inflammation and pneumonia by regulating the TLR4/NF-kB signaling pathway. Journal of Translational Medicine 20, 130 (2022).

48. Sun, X. et al. Anti-inflammatory mechanisms of the novel cytokine interleukin-38 in allergic asthma. Cellular & molecular immunology 17, 631–646 (2020).

49. Pan, W. et al. Identification of Potential Differentially-Methylated/Expressed Genes in Chronic Obstructive Pulmonary Disease. Copd 20, 44–54 (2023).

50. Headland, S.E. et al. Oncostatin M expression induced by bacterial triggers drives airway inflammatory and mucus secretion in severe asthma. Sci Transl Med 14 (2022).

51. Chiba, Y. et al. Altered renin-angiotensin system gene expression in airways of antigen-challenged mice: ACE2 downregulation and unexpected increase in angiotensin 1–7. Respiratory Physiology & Neurobiology 316, 104137 (2023).

52. Bouma, F., Nyberg, F., Olin, A.C. & Carlsen, H.K. Genetic susceptibility to airway inflammation and exposure to short-term outdoor air pollution. Environ Health-Glob 22 (2023).

53. Silveyra, P. & Floros, J. Genetic complexity of the human surfactant-associated proteins SP-A1 and SP-A2. Gene 531, 126–132 (2013).

54. Blanchet, M.R. et al. CD34 facilitates the development of allergic asthma. Blood 110, 2005–2012 (2007).

55. Cheng, Y.Q. et al. Elucidation of the mechanisms and molecular targets of KeChuanLiuWei-Mixture for treatment of severe asthma based on network pharmacology. Chem Biol Drug Des (2023).

56. Chen, Y.J. et al. Inactivation of the tumor suppressor p53 by long noncoding RNA RMRP. P Natl Acad Sci USA 118 (2021).

57. Xing, Y. et al. Comprehensive analysis of differential expression profiles of mRNAs and lncRNAs and identification of a 14-lncRNA prognostic signature for patients with colon adenocarcinoma. Oncology reports 39, 2365–2375 (2018).

58. Li, D. et al. Long noncoding RNA FGD5-AS1 promotes colorectal cancer cell proliferation, migration, and invasion through upregulating CDCA7 via sponging miR-302e. In Vitro Cell Dev-An 55, 577–585 (2019).

59. Yang, F., Chen, L. & Wang, Z.J. MicroRNA-32 inhibits the proliferation, migration and invasion of human colon cancer cell lines by targeting E2F transcription factor 5. Eur Rev Med Pharmaco 23, 4156–4163 (2019).

60. Guan, Y.F. et al. Upregulation of HSPA1A/HSPA1B/HSPA7 and Downregulation of HSPA9 Were Related to Poor Survival in Colon Cancer. Front Oncol 11 (2021).

61. Belaguli, N.S. et al. GATA6 Promotes Colon Cancer Cell Invasion by Regulating Urokinase Plasminogen Activator Gene Expression. Neoplasia 12, 856–U814 (2010).

62. Lai, H.T., Chiang, C.T., Tseng, W.K., Chao, T.C. & Su, Y. GATA6 enhances the stemness of human colon cancer cells by creating a metabolic symbiosis through upregulating expression. Mol Oncol 14, 1327–1347 (2020).

63. Bhatlekar, S., Viswanathan, V., Fields, J.Z. & Boman, B.M. Overexpression of HOXA4 and HOXA9 genes promotes self-renewal and contributes to colon cancer stem cell overpopulation. J Cell Physiol 233, 727–735 (2018).

64. Yu, Y.F. et al. The inhibitory effects of COL1A2 on colorectal cancer cell proliferation, migration, and invasion. J Cancer 9, 2953–2962 (2018).

65. Chen, J. et al. Type-2 11β-hydroxysteroid dehydrogenase promotes the metastasis of colorectal cancer via the Fgfbp1-AKT pathway. Am J Cancer Res 10, 662-+ (2020).

66. Liu, Z.-P., Wu, C., Miao, H. & Wu, H. RegNetwork: an integrated database of transcriptional and post-transcriptional regulatory networks in human and mouse. Database 2015, bav095 (2015).

67. Han, H. et al. TRRUST: a reference database of human transcriptional regulatory interactions. Scientific reports 5, 11432 (2015).

68. Bovolenta, L., Acencio, M. & Lemke, N. HTRIdb: an open-access database for experimentally verified human transcriptional regulation interactions. Nature Precedings, 1–1 (2012).

69. Van Landeghem, S. et al. Exploring biomolecular literature with EVEX: connecting genes through events, homology, and indirect associations. Advances in bioinformatics 2012 (2012).

70. Mathelier, A. et al. JASPAR 2014: an extensively expanded and updated open-access database of transcription factor binding profiles. Nucleic acids research 42, D142–D147 (2014).

71. Lachmann, A. et al. ChEA: transcription factor regulation inferred from integrating genome-wide ChIP-X experiments. Bioinformatics 26, 2438–2444 (2010).

72. Matys, V. et al. TRANSFAC® and its module TRANSCompel®: transcriptional gene regulation in eukaryotes. Nucleic acids research 34, D108–D110 (2006).

73. Xie, X., Rigor, P. & Baldi, P. MotifMap: a human genome-wide map of candidate regulatory motif sites. Bioinformatics 25, 167–174 (2009).

74. Feingold, E. et al. The ENCODE (ENCyclopedia of DNA elements) project. Science 306, 636–640 (2004).

75. Kamburov, A., Stelzl, U., Lehrach, H. & Herwig, R. The ConsensusPathDB interaction database: 2013 update. Nucleic acids research 41, D793–D800 (2013).

76. Szklarczyk, D. et al. STRING v11: protein–protein association networks with increased coverage, supporting functional discovery in genome-wide experimental datasets. Nucleic acids research 47, D607–D613 (2019).

77. Razick, S., Magklaras, G. & Donaldson, I.M. iRefIndex: a consolidated protein interaction database with provenance. BMC bioinformatics 9, 1–19 (2008).

78. Huang, J.K. et al. Systematic evaluation of molecular networks for discovery of disease genes. Cell systems 6, 484–495. e485 (2018).

